# Comprehensive Analysis of Regenerative and Transformed Liver Reveals Distinct, Early Metabolic Alterations in Cancer

**DOI:** 10.1101/2022.09.10.507425

**Authors:** Daniel J. McLaughlin, Hannan Mir, Minwoo Nam, Richard Possemato

## Abstract

Alterations in cellular metabolism represent an important response to proliferative signals in both normal and transformed tissues. The benign proliferative process of liver regeneration after partial hepatectomy offers insight into homeostatic mechanisms to control liver mass, which are disrupted in liver disease induced by viral factors, alcohol, or associated with obesity. Moreover, successful targeting of cancer depends on the identification of genes and pathways that are selectively activated in the transformed state. Here, we present a differential transcriptomic and metabolomic analysis of benign proliferative and transformed liver, including associated plasma metabolite and lipid species. Using partial hepatectomy-induced liver regeneration and diethylnitrosamine (DEN) induced carcinogenesis, we identify and analyze alterations specific to multiple regenerative and transformed tissue states. Transcriptomics and LC/MS based metabolite profiling reveal fatty acid import and storage are specifically rewired during liver regeneration in a time dependent manner, a phenomenon not observed in liver tumors. In contrast, liver tumors exhibit preferential activation of numerous metabolic pathways, including glycolysis, serine biosynthesis, and polyamine metabolism. Alterations in serine metabolism occur at the earliest detectable stages in tumorigenesis and promote survival upon serine restriction. These data demonstrate that transformation-induced alterations in metabolism are distinct from those observed in normal regenerative cell division, which may be used to identify transformation-specific liabilities.

## Introduction

The mouse liver is an intriguing organ to ascertain how activation and suppression of metabolic pathways supports proliferation and transformation because of its regenerative capacity following injury and the availability of chemical carcinogens to induce liver tumors. Initiation of liver regeneration after partial hepatectomy induces a synchronous entry into the cell cycle for most hepatocytes that compose the organ.^1^ Partial hepatectomy, or the surgical resection of one or multiple lobes of the liver, provides the necessary stimulus to induce hepatocyte re-entry into the cell cycle from their initial quiescent state.^1^ Hepatocytes are the first cells within the liver to undergo cell cycle re-entry, and each cell undergoes 1-2 rounds of replication before the original liver mass is restored.^1^ After partial hepatectomy, liver mass is restored exclusively through hepatocyte proliferation, and not through the activation of hepatic progenitor cells.^2^

Peritoneal treatment of diethylnitrosamine (DEN) in mice is an established model for studying the alterations that underlie liver cancer formation. Upon administration, DEN is rapidly metabolized by the liver specific cytochrome P450 enzymes including CYP2E1.^3^ The product of CYP2E1 mediated DEN dealkylation is ethyldiazohydroxide, which is further decomposed to produce a set of highly reactive and transient electrophiles that ethylate macromolecules throughout the cell. Guanine is commonly ethylated in this fashion leading to the formation of the DNA adducts N7-ethylguanine and O6-ethylguanine, which often precipitates guanine to thymine transversions or direct base mispairing.^4^ Tumors generated in this manner contain numerous somatic single nucleotide variants, often in Hras, Braf, or Egfr.^5^ The mutational burden in DEN tumors makes this model particularly interesting. Other models use genetically engineered mice to induce tumorigenesis that carry deletions or point mutations in only a small set of genes.^6^ This lack of tumor mutations differs greatly from tumors found in human HCC, which often have a high tumor mutational burden. Indeed, DEN induced tumors share high similarity to a subset of human HCC with RAS/MAPK pathway mutation and poor prognosis.^5^ Given the ease through which we can chemically induce tumor formation in the mouse liver, we are able to efficiently compare rapidly growing liver tumors to the rapid growth that occurs within the same tissue after hepatectomy.

Here, we harvest regenerating liver tissue or liver tumors for analysis of steady state metabolism and metabolite utilization using RNAseq and LC/MS-based measurements of metabolite levels. We perform a comprehensive metabolic analysis of liver tissue at several time points during the regenerative process and control for differences related to diurnal fluctuations in metabolism. These experiments provide the opportunity to assess, in vivo, key differences in the metabolism of quiescent, proliferative, and transformed tissues.

## Results

### Partial Hepatectomy and Carcinogen Induced Transformation Models

To induce liver regeneration, we performed partial hepatectomy on a cohort of adult male mice. For each mouse, we collected an initial, quiescent liver sample or plasma followed by a terminal collection of regenerating tissue or plasma at 12, 24, 36, 48, 72, or 96 hours post hepatectomy (Supplementary Figure 1A and 1B). Of note, the regenerating liver sample collected is a separate lobe from that which has been surgically removed. To account for changes in gene expression that are not associated with regeneration (e.g. diurnal associated shifts in expression or inflammatory signaling associated with surgery), we performed a set of sham surgeries on a separate cohort of mice. An incision was made into the peritoneal cavity of each mouse, and a sterile cotton swab was used to palpate the liver, but no tissue was removed. We then collected terminal liver samples or plasma from these sham mice at the same time points as those of the regeneration group.

In a subset of mice, we administered an intraperitoneal dose of BrdU 2 hours before euthanasia. BrdU is a thymidine analogue incorporated into dividing DNA during S phase. To verify that the livers of the regeneration group have begun regeneration, we performed histological staining for BrdU at each time point of regeneration (Supplementary Figure 1C). We observed maximum BrdU incorporation in the regenerating liver 48 hours post hepatectomy followed by a steep drop-off in incorporation at 72 hours indicating that we were successful in activating a regenerative response, and that the hepatocytes that make up the liver reach S phase by 48 hours post hepatectomy.

To identify metabolic pathways rewired in hepatic tumors, we treated a separate cohort of weaning male mice with the hepatocarcinogen diethylnitrosamine (DEN) at a dose of 0.025 mg/g body weight (Supplementary Figure 1D). Male mice exhibit higher sex associated expression of several cytochrome p450 oxidase genes involved in xenobiotic compound metabolism and are therefore more susceptible to DEN toxicity.^4^ Histological staining revealed that visible lesions appear by 20 weeks in male mice, which progress to larger tumors at 30 weeks post DEN treatment (Supplementary Figure 1E). After 30 weeks, we harvested 18 tumors in total from the adult mice.

### Diurnal Shifts in Liver Metabolism

Because diurnal changes in metabolism could obfuscate changes related to tissue regeneration, we first compared samples collected following sham surgery. We performed global transcriptomic and untargeted LC/MS based metabolomic analyses using the collected tissue samples and plasma and performed hierarchical clustering of gene transcripts and metabolites significantly altered (Supplementary Tables 1-6). We generated a heatmap with hierarchical clustering containing these genes and performed a gene ontology analysis on each gene cluster (Figure 1A). In this analysis, we implemented the DAVID suite of bioinformatic resources to identify relevant biological processes represented in each gene cluster (Figure 1B).^7^ We chose biological process and gene ontology data sets that contained a minimum of 2 genes from out data set and achieved a false discovery rate below 0.5 (FDR < 5.00E-1) for further analysis. We predicted that alterations in these genes and metabolites are the result of diurnal rhythms rather than the stimulus from partial hepatectomy.

**Figure 1.**
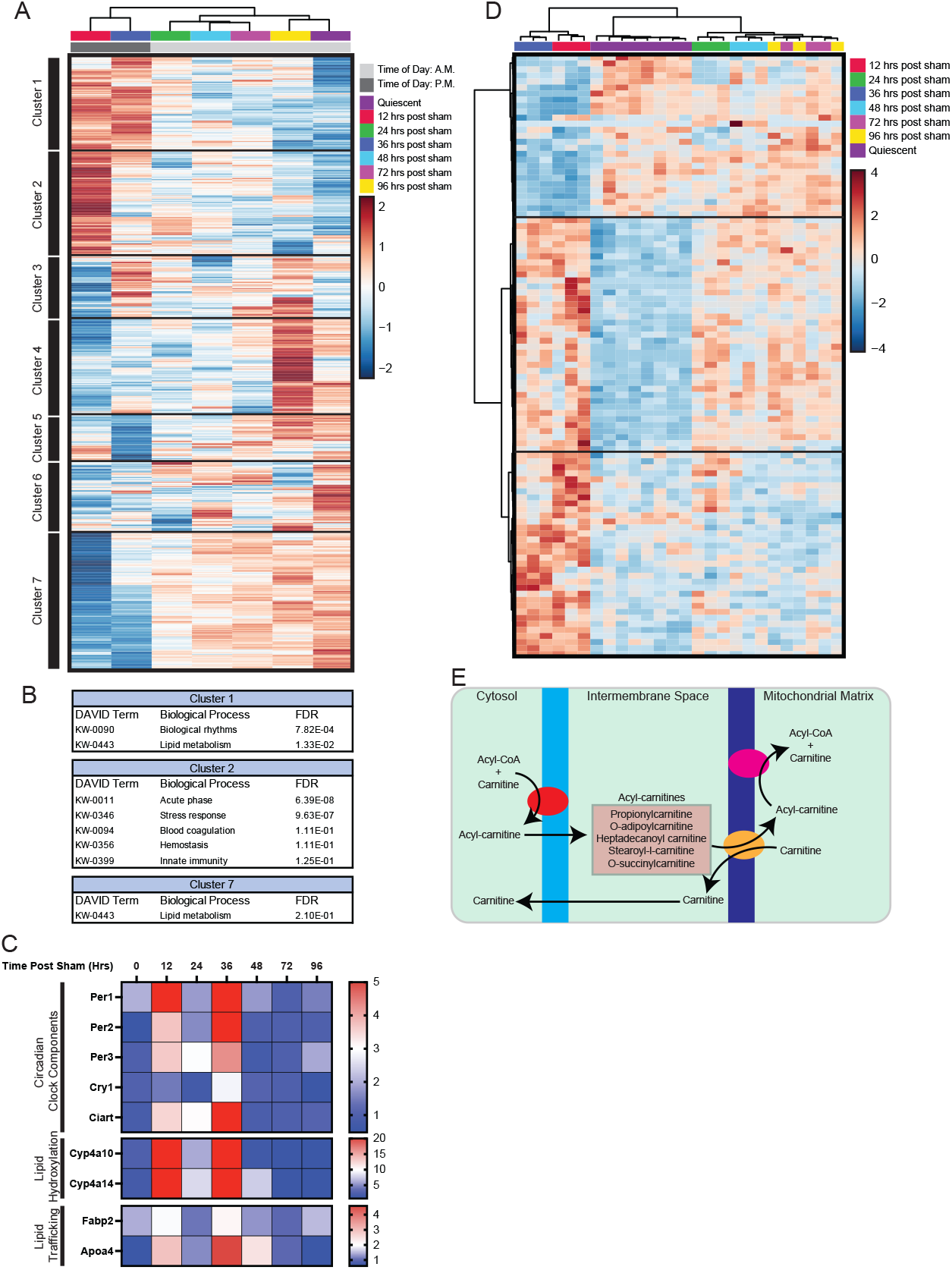
Significantly altered transcripts and metabolites in animals undergoing sham surgery. A, Hierarchically clustered heatmap of top 1000 significantly altered transcripts in liver tissue after sham surgery. Log_2_ CPM is reported in the heatmap. Gene order reported in Supplementary Table 7. B, Analysis of genes enriched in the indicated clusters from *A*, using the DAVID suite of software. Terms listed that contain at least 2 genes and an FDR <0.5 (Supplementary Table 7). C, Data from *A* for the specific genes indicated, relative to 72 hr time point. D, Hierarchically clustered heatmap of top 100 significantly altered metabolites in liver tissue after sham surgery. Data are row normalized and the Log_2_ relative abundance is reported in the heatmap. Metabolite order reported in Supplementary Table 7. E, Schematic of the carnitine shuttle.

We identified 7 unique gene clusters from our hierarchical analysis of transcripts after sham surgery (Figure 1B and Supplementary Table 7). The time points at night, 12 and 36 hours post sham surgery, formed a distinct expression cluster from the daytime samples. Sham Gene Cluster 1 and 7 contains genes upregulated and downregulated at night, respectively and fall into two main pathways: biological rhythms and lipid metabolism (Figure 1C). The period circadian clock genes Per1, 2, and 3 are upregulated at these times as well as Nocturnin (Noct), cryptochrome 1 (Cry1), and circadian associated repressor of transcription (Ciart). Per1, Per2, Per3, Cry1, and Ciart encode proteins that act as some of the core components of the internal time-keeping system within mammals known as the circadian clock, which regulates several metabolic processes. One of these processes is lipid metabolism. We observed elevation in the genes encoding lipid trafficking enzymes Fabp2 and Apoa4. This pathway may be activated as a response to increased activity and feeding that takes place during this period. In support of this notion, we observed elevation in the cytochrome p450 genes Cyp4a10 and Cyp4a14, which encode peroxisomal enzymes that catalyze very long chain fatty acid oxidation after feeding and are targets of regulation via the circadian clock.^8^

Metabolomic analysis revealed changes in lipid metabolism concordant with the diurnal clustering observed in our transcriptomic analysis (Figure 1D and Supplementary Table 7). Sham Metabolite Cluster 3 contains a set of metabolites that are more abundant at 12 and 36 hours post sham compared to all other periods, and includes carnitine and the acylcarnitines propionylcarnitine, o-adipoylcarnitine, heptadecanoyl carnitine, stearoyl-l-carnitine, and o-succinylcarnitine. Fatty acids, particularly long chain fatty acids, are unable to defuse through the mitochondrial membrane without the help of the carnitine transport system. The enzyme CPT1 converts carnitine and acyl-CoAs to acylcarnitines as they are passed through the outer mitochondrial matrix. These acylcarnities are then converted back to acyl-CoAs and carnitine and deposited into the mitochondrial matrix for subsequent oxidation reactions (Figure 1E).^9^ Thus, acylcarnitines like the ones in Sham Metabolite Cluster 3 are generally regarded as markers for increased fatty acid uptake and flux into the mitochondria, indicating an increase in fatty acid trafficking during the night periods. Furthermore, recent research has shown that the liver experiences an increase in acylcarnitines directly after feeding in both mice and humans.^10^

We also observed changes in gene expression that do not correlate with time of day. These genes are contained in Sham Gene Clusters 2-6 (Figure 1A). We predict that these changes are not the result of diurnal regulation but rather a response to the sham surgery. We observed acute phase inflammation, stress response, and immunity as the top enriched gene sets. Similarly, some metabolites exhibit elevated abundance at all times post sham. These metabolites may also be associated with inflammation.

### Metabolic Shifts in Liver Metabolism after Partial Hepatectomy

To determine which pathways are altered exclusively by regeneration, we calculated the percent difference between the transcript abundance of each time post hepatectomy compared to its corresponding sham samples. We then selected transcripts whose abundance changed by greater than 50% compared to the sham samples to select for transcripts selectively altered during regeneration (Figure 2A, Supplementary Table 7).

**Figure 2.**
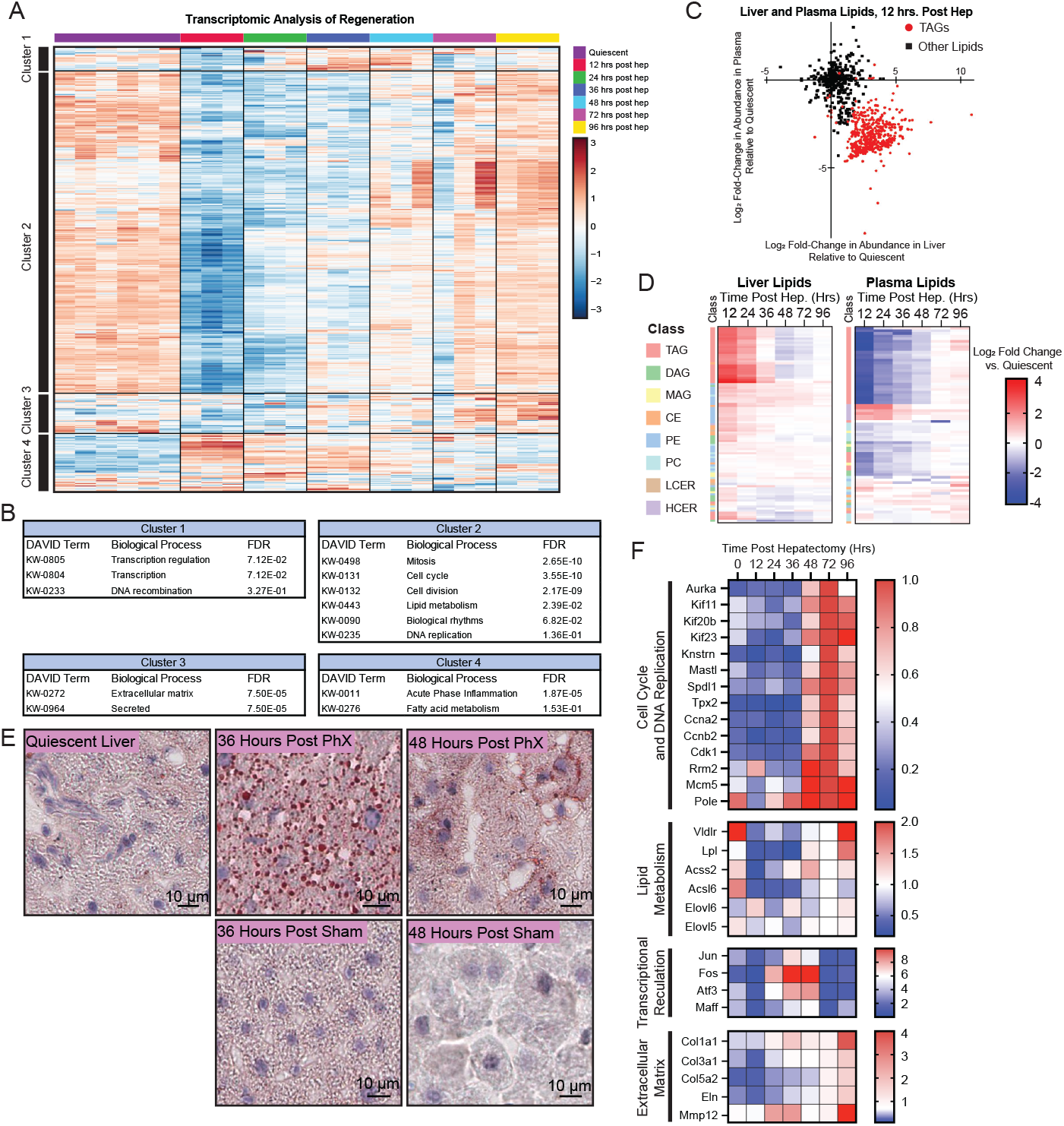
Significantly altered transcripts and metabolites during liver regeneration. A, Hierarchically clustered heatmap of top 500 significantly altered transcripts from liver tissue at the indicated time points after partial hepatectomy (quiescent, n=6; 12-96 hr. regeneration time points, n=3). Log_**2**_ CPM is reported in the heatmap. Gene order reported in Supplementary Table 7. B, Analysis of genes enriched in the indicated clusters from *A*, using the DAVID suite of software. Terms listed that contain at least 2 genes and an FDR <0.5 (Supplementary Table 7). C, Relative level of lipids (Log_2_ fold change compared to quiescent) at 12 hrs after partial hepatectomy, in the plasma versus liver tissue. Triglyceride species (TAGs) are red circles, other lipids are black squares. D, Hierarchically clustered heatmap of significantly altered lipids from liver tissue or plasma at the indicated time point after partial hepatectomy (p < 0.05, quiescent, n=8; 12-96 hr. regeneration time points, n=6). Log_2_ fold change relative to quiescent reported. Lipid class indicated by colored bar to the left of each heatmap. Lipid order reported in Supplementary Table 7. E, Histological staining for neutral lipids (Oil Red O) on representative frozen sections collected at the indicated time point after hepatectomy, eosin counter-stain. F, Data from *A* for the specific genes indicated, relative to 72 hr time point.

We identified 4 unique gene clusters in our transcriptomic analysis of partial hepatectomy treated samples (Figure 2A, 2B, Supplementary Table 7). Of note, in Regenerating Gene Cluster 4, we identified acute phase inflammation as a major pathway activated during the initial twelve hours post hepatectomy. We also observed a down regulation of cell cycle related gene sets seen in Regenerating Gene Cluster 2 in the first thirty-six hours of regeneration followed by a rebound and eventual upregulation starting forty-eight hours post hepatectomy. These genes include aurora kinase A (Aurka), mitosis associated genes Kif11, Kif20b, Kif23, Knstrn, Mastl, Spdl1, Tpx2, cyclins Ccna2 and Ccnb2, as well as the cyclin dependent kinase Cdk1. Additionally, the gene that encodes the ribonucleotide reductase Rrm2, which catalyzes the synthesis of deoxyribonucleotides required for DNA synthesis, peaks in expression 48 hours after hepatectomy. The helicase encoding gene Mcm5 and the DNA polymerase encoding gene Pole also peak in expression at this time, consistent with DNA replication occurring in accordance with our BrdU incorporation data.

We also observed substantial fluctuations in metabolites and transcripts associated with lipid metabolism. We observed a peak in major membrane lipid species (phosphatidylcholine and phosphatidylethanolamine) and cholesterol esters at 12 hours after partial hepatectomy, falling to baseline by 96 hours (Figure 2C-D, Supplementary Table 7). Triacylglycerol (TAG) species were also broadly elevated 12-24 hours after partial hepatectomy, falling below baseline levels at 48-72 hours, before returning to baseline at 96 hours. These changes in liver TAG species was mirrored by a corresponding decrease in these same lipids from the plasma at 12-24 hours, consistent with either increased absorption of TAGs by the liver or decreased release from the liver at the onset of regeneration (Figure 2C-D). A neutral lipid stain revealed the transient appearance of lipid droplets within the liver, frequently observed in partial hepatectomy models of liver regeneration,^11^ at 36 hours post hepatectomy, a phenomenon absent from the sham samples, diminishing in later time points (Figure 2E). This peak in TAG levels was followed by a peak in lipid metabolism-related gene expression starting 36 hours post hepatectomy (Figure 3F). These genes include the very low-density lipoprotein receptor (Vldlr), lipoprotein lipase (Lpl), short and long chain acyl-CoA synthetases (Acss2 and Acsl6), and fatty acid lengthening enzyme 5 and 6 (Elovl5 and Elovl6). Elovl5 and 6 encode enzymes associated with adipogenesis and increased intracellular triglyceride abundance.^12^ Together, this gene expression profile indicates extracellular fatty acids as an important factor in the regenerating liver’s metabolism. The protein VLDLR facilitates import of triglyceride rich VLDL particles from the blood supply. Membrane bound LPL hydrolyzes these triglycerides and releases free fatty acids into the cell.^13^ The acyl-CoA synthetases convert these fatty acids to fatty acid acyl-CoA molecules which fulfill several functions in the cell. Elovl3 catalyzes the elongation of fatty acid acyl-CoAs to produce very long chain fatty acid acyl-CoAs. These long fatty acids can be oxidized to provide energy, serve as intermediates for the synthesis of other lipids, or they can fuel triglyceride accumulation.^14^ Other studies have shown that ablation of Elovl3 gene expression leads to impaired lipid storage in mice.^12^ Taken together, these data demonstrate that TAGs are a dynamic intermediate during liver regeneration, possibly to fuel ATP and membrane lipid synthesis.

**Figure 3.**
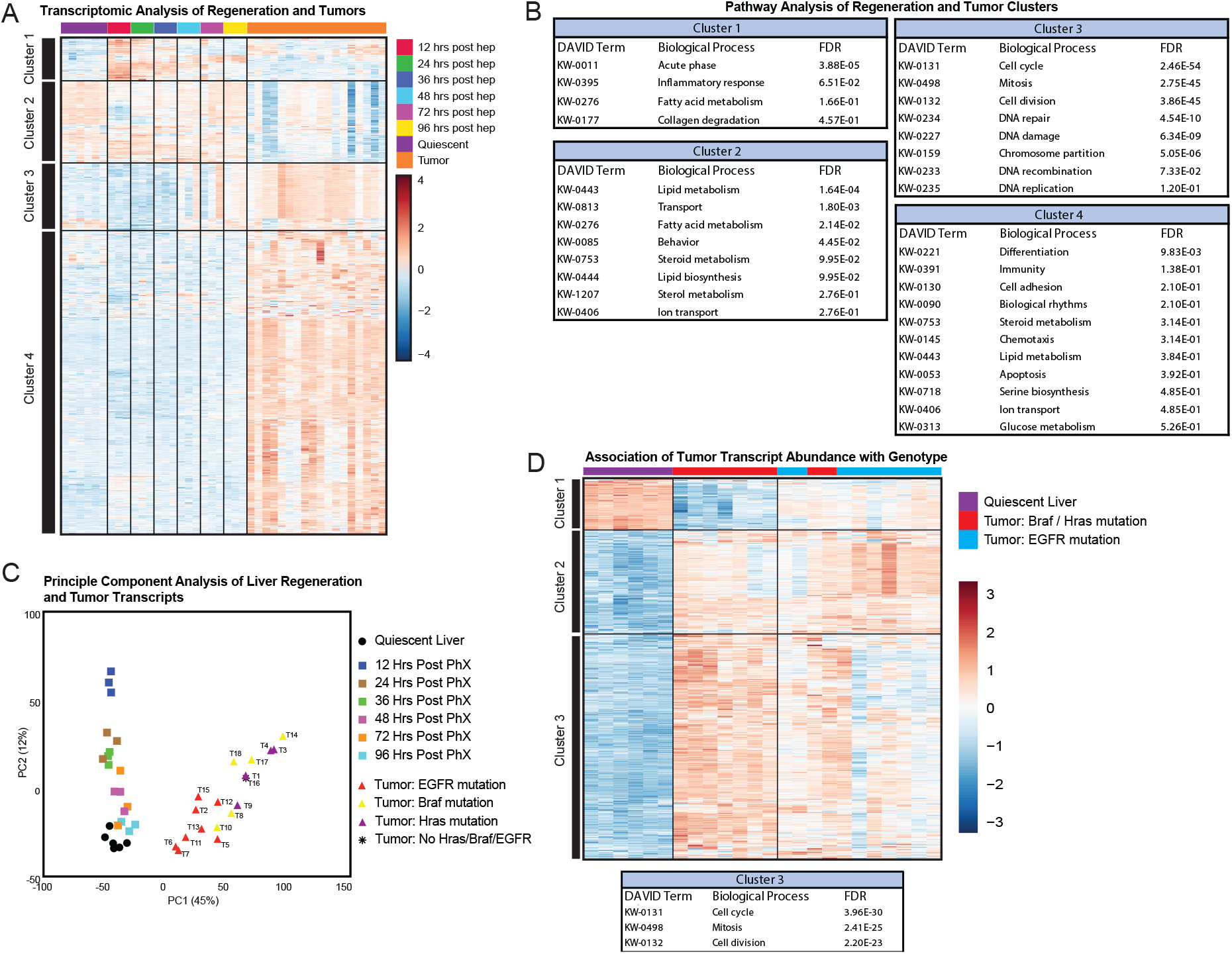
Transcripts significantly altered in liver tumors compared to regenerating and quiescent liver. A, Hierarchically clustered heatmap of top 1000 significantly altered transcripts in liver tumors or regenerating liver (tumors, n=18, quiescent, n=6; 12-96 hr. regeneration time points, n=3). Log_**2**_ CPM is reported in the heatmap. Gene order reported in Supplementary Table 7. B, Analysis of genes enriched in the indicated clusters from *A*, using the DAVID suite of software. Terms listed that contain at least 2 genes and an FDR <0.5 (Supplementary Table 7). C, Principle Component Analysis of data from *A*. For tumors, the Hras Q61R/L, Braf V585E, or Egfr F254I mutation status is indicated. D, Hierarchically clustered heatmap of top 1000 significantly altered transcripts in liver tumors versus quiescent liver tissue (tumors, n=18, quiescent, n=6). Log_**2**_ CPM is reported in the heatmap. Hras, Braf, and Egfr mutation status indicated. Below, analysis of genes enriched in the indicated cluster, using the DAVID suite of software. Terms listed that contain at least 2 genes and an FDR < 1e-5 (Supplementary Table 7).

Starting 24 hours post hepatectomy and peaking at 36 hours, we noticed a set of transcriptional regulators in the AP-1 family increase in expression seen in Regenerating Gene Cluster 1 (Figure 2F). This family includes Jun, Fos, Atf3, and Maff. The proteins encoded by these members form dimers that bind to DNA and regulate transcription. AP-1 family members have been shown to enhance cell cycle progression by regulating the expression of the enzymes required for DNA synthesis and several cyclin genes including Ccna2 and Ccnb2. We surmise that this AP-1 pathway is responsible for the progression into S phase that occurs shortly after these genes peak in expression.

Regenerating Gene Cluster 3 contains a gene set composed of several components of the extracellular matrix that also change in expression in a pattern similar to the cell cycle related gene sets mentioned previously (Figure 2F). Collagen type 1, 3, and 5 (Col1a1, 3a1, and 5a2), elastin (Eln), and matrix metallopeptidase 12 (Mmp12) are immediately suppressed during regeneration but rise in expression over 96 hours. This is indicative of the robust tissue remodeling that occurs during the process of regeneration.

### Tumor Specific Alterations in DEN Induced Tumors

We hypothesized that proliferation and cell cycle progression related genes would represent the primary similarity between regenerating and transformed tissue. To test this, we performed hierarchical clustering and pathway analysis on the combined data set that included quiescent, regenerating, and transformed tissue (Figure 3A-3B, Supplementary Table 7). From this analysis, we identified 4 distinct gene clusters (Figure 3B). Combined Gene Cluster 3 from this analysis represents the genes that are upregulated both during regeneration and in tumor growth. The genes in this cluster mapped to several different biological processes including genes required for DNA replication, cell cycle progression, or cell division, indicating that the same cell cycle progression and proliferation related genes are enhanced in both states. These genes, therefore, represent key pathways that are required for both the normal and transformed proliferation of liver cells.

Tumors strongly clustered together by gene expression, and principle component analysis revealed that PC1 delineated transformed versus non-transformed samples (Figure 3C). PC2 differentiated regeneration samples from each other in a time-dependent manner, with the 12 hour regeneration samples exhibiting the greatest difference, which diminished in subsequent samples. DNA collected from tumors was used to identify the mutation status of oncogenic driver known to be present in DEN-induced tumors (Hras Q61R/L, Braf V585E, Egfr F254I) (Supplementary Table 8). Interestingly, the distance along both PC1 and PC2 axes also correlated with oncogenic driver mutation, with Hras- and Braf-mutated tumors clustering together, further away from quiescent tissue. Indeed, genes related to cell cycle and cell division were most strongly upregulated in Hras- and Braf-mutated tumors, compared to Egfr-mutated tumors, while those related to liver specific function were more suppressed, indicative of a less differentiated tumor (Figure 3D). Among the genes most upregulated in Hras- and Braf-mutated tumors compared to Egfr-mutated tumors are the immune checkpoint receptor Cd40, CDK inhibitors Cdk2na and Cdkn2b, and stress response gene Lcn2. Of these, Lcn2 was also very strongly induced during regeneration.

We also performed pathway analysis of Combined Gene Cluster 4, which represents genes up regulated in DEN tumors but not during regeneration (Figure 3B). This list of tumor specific genes proved to be more diverse than the list of up-regulated genes common to both tissue states. We identified several metabolic processes specific to DEN tumors. We noted changes in gene expression and metabolite abundance indicating altered glucose metabolism, altered polyamine metabolism, and enhanced serine biosynthesis, as described in detail below.

### Glycolysis Side-Branches are Altered in DEN Induced Tumors

Several metabolites are more abundant in DEN induced tumors compared to regenerating tissue (Supplementary Table 3 and Supplementary Figures 2-3). First, we noticed an increase in concentrations of the glucose derivatives glucose-6-phosphate, fructose-6-phsophate, 1,5-anhydro-D-glucitol, and glucosamine-6-phosphate (Figure 4A). Most of the glucose that enters a cell is quickly converted to glucose-6-phosphate by hexokinase, which is a major node in both glycolysis and the pentose phosphate pathway. Glucose-6-phosphate is isomerized to fructose-6-phosphate to fuel glycolysis or dehydrogenated to 6-phosphogluconate to fuel the pentose phosphate pathway. Fructose-6-phosphate and 6-phosphogluconate are both elevated in DEN tumors, indicating flux through glycolysis as well as its side-branches like the pentose phosphate pathway. These tumors exhibit elevated expression of the gene pyruvate dehydrogenase kinase 3 (Pdk3), an inhibitor of the pyruvate dehydrogenase complex. The enzyme PDK3 reduces pyruvate entry into the mitochondrion and has been shown to stimulate compensatory flux through glycolysis. Overexpression of Pdk3 has been noted as a marker for poor prognosis in many cancer types including melanoma, glioma, gastric cancer and ovarian cancer.^15^ Furthermore, we observed upregulation in glucose-6-phosphate dehydrogenase (G6pdx), which encodes an enzyme that catalyzes the initial glucose-6-phosphate dehydrogenation in the pentose phosphate pathway (Figure 4B).

**Figure 4.**
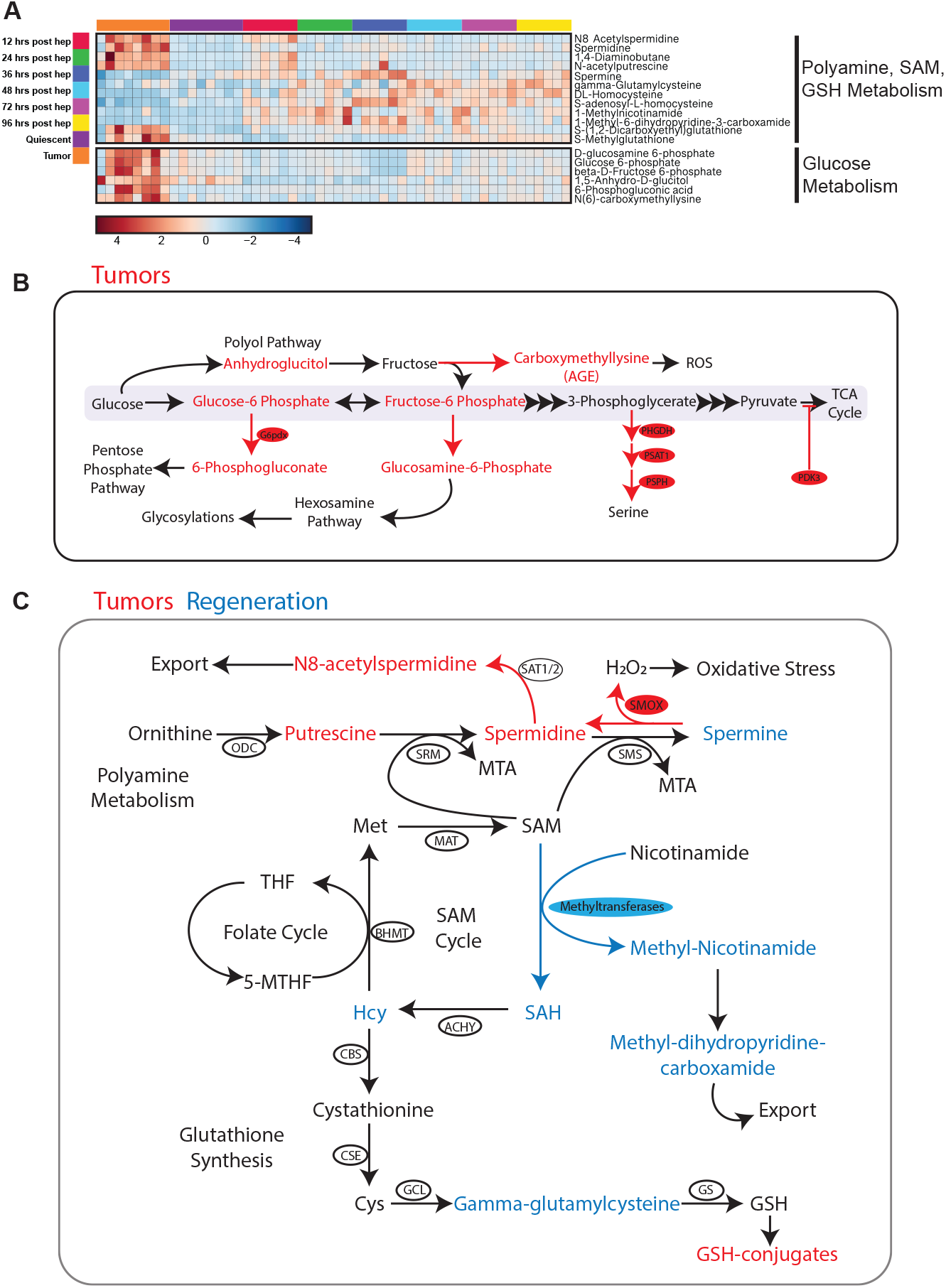
Metabolites significantly altered in liver tumors compared to regenerating and quiescent liver. A, Selected significantly altered metabolites in liver tumors, quiescent liver, or regenerating liver. Relative log_**2**_ fold change is reported in the heatmap. B-C, Schematic of relevant metabolic pathways with significantly altered enzymes and metabolites indicated.

1,5-anhydro-D-glucitol, also known as sorbitol, is a product of the polyol pathway, another glycolysis side-branch where glucose is reduced by aldose reductase consuming NADPH as a cofactor.^16^ High concentrations of intracellular glucose activate aldose reductase to stimulate sorbitol production.^16^ In diabetes and nonalcoholic fatty liver disease, activation of this pathway is stimulated by robust glucose uptake in the liver and is generally predicted to exacerbate oxidative stress within the cell caused by a depletion in reduced glutathione as a result of NADPH depletion.^16^ Advanced glycosylation end products (AGEs) are heavily associated with the oxidative stress caused by hyperglycemia in these diseases.^17^ Indeed the abundance of N(6)-carboxymethyllysine, a common marker for AGEs, is highly induced in these tumors compared to healthy dividing tissue (Figure 4A-B). We also noted an increase in the glutathione conjugates S-(1,2-Dicarboxyethyl) glutathione and S-methylglutathione. Accumulation of these specific metabolites indicates a cellular response to oxidative stress and the active utilization of glutathione.^18^

Glucosamine-6-phosphate is an intermediate in the hexosamine biosynthesis pathway, a side-branch of glycolysis responsible for the production of glycosylation substrates while consuming fructose-6-phosphate and glutamine.^19^ The hexosamine biosynthesis pathway products are used to form glycolipids, proteoglycans, and glycosaminoglycans. Tumors often exhibit aberrant glycosylation, specifically in melanoma where activation of the hexosamine biosynthesis pathway regulates cell migration and metastasis.^19^ The intermediates of these glycolytic side-branches are not upregulated at any time during regeneration.

### Polyamine Metabolism is Altered in DEN Induced Tumors

In DEN induced tumors we observed increases in the intermediates of polyamine metabolism, including N-acetylputrescine, N8-acetylspermidine, 1,4-diaminobutane, and spermidine (Figure 4A and 4C). Interestingly, spermine, another polyamine, was not elevated in DEN induced tumors, but was instead elevated in regenerating tissue 36 hours post hepatectomy. The polyamine pathway begins with the formation of 1,4-diaminobutane (putrescine) by the enzyme ornithine decarboxylase. Through a series of irreversible aminopropyl group additions, putrescine is converted into spermidine and spermine by spermidine synthase and spermine synthase respectively (Figure 4A and 4C).^20^ Spermine can be converted back to spermidine through a separate oxidation reaction catalyzed by spermine oxidase (SMOX), which produces hydrogen peroxide in the process. Additionally, in a reaction catalyzed by spermidine/spermine N1-acetyltransferase 1 (SSAT), spermine is converted to N-acetylspermine, which is subsequently exported from the cell.^20^ In addition to the increased concentration of these polyamine metabolites, we observed an increase in the expression of Smox. We predict the decrease in spermine within these DEN induced tumors is the result of SMOX activity, which fuels its catabolism. Upregulation in Smox transcription is sufficient to produce oxidative stress *in vitro* due primarily to the hydrogen peroxide produced in this reaction.^21^ Further, human hepatocellular carcinoma often exhibits upregulated SMOX gene expression as well as increased spermine metabolism. SMOX high tumors have worse prognosis and faster growth rates compared to tumors with normal SMOX gene expression.^22^ Given the reduced spermine concentration in DEN induced tumors compared to regenerating tissue, we hypothesize that these tumors preferentially catabolized spermine to make spermidine and other polyamines, while the regenerating tissue preferentially synthesizes spermine.

### The SAM Cycle May be More Active in Regenerating Tissue Than in DEN Induced Tumors

In our combined metabolomics data we observed an increase in the abundance of several SAM cycle intermediates during regeneration and not in DEN induced tumors (Figure 4A and 4C). S-adenosylmethionine (SAM) is a metabolite central to several metabolic processes including methylation reactions, one-carbon metabolism, autophagy, and polyamine synthesis. SAM is the primary methyl doner for DNA, RNA, lipid, and histone methylation.^23^ This methylation can regulate gene transcription and protein activity. SAM is synthesized normally in hepatocytes and is mainly derived from dietary methionine. The methyltransferase reactions required for methylation consume SAM and produce S-adenosylhomocysteine (SAH). SAH is subsequently converted via the transsulfuration pathway into homocysteine, cystathionine, and cysteine, a critical component of glutathione, or can be recycled to SAM by the one carbon pool ^23^ (Figure 4A and 4C). Our metabolic profiling indicates an increase in SAH and homocysteine abundance 36 hours post hepatectomy. Because SAH is produced as a result of SAM mediated methylation reactions, the increased abundance in these metabolites indicates robust methylation in regenerating tissue. This increase in methylation may also be linked to the observed changes in gene expression and lipid metabolism post-hepatectomy. We also observe an increase in plasma formiminoglutamate (FIGLU), particularly at 12-36 hours after partial hepatectomy (Supplementary Figure 4). FIGLU is a catabolite of histidine and a marker for defects in the one carbon pool. Thus, stress placed on the one carbon pool because of increased nucleotide synthesis may be resulting in catabolism of SAH to homocysteine, as one carbon units needed to drive recycling of SAH to methionine are scarce.

We also observed an increase in gamma-glutamylcysteine, an intermediate in the synthesis of glutathione, 48 hours post hepatectomy. Thus, increased transsulfuration may be fueling glutathione biosynthesis 36-48 hours post hepatectomy, although we did not detect changes in reduced glutathione. The increased SAM cycle flux may also explain the difference in spermine concentrations between DEN tumors and regenerating tissue given that SAM is a necessary cofactor for the production of spermine.^20^ Spermine peaks in concentration 36 hours post hepatectomy, at the same time we observe peak abundance of other SAM cycle metabolites. DEN induced tumors show enhanced oxidation of spermine while regenerating tissue show enhanced synthesis of spermine and glutathione. Together, these data indicate that the SAM cycle flux, preferentially activated during regeneration, may lead to epigenetic modifications and redox control mechanisms not active in tumors.

### Serine Biosynthesis Enzymes are Upregulated in DEN Induced Tumors

Because we noted strong over-expression of the serine biosynthesis pathway enzymes in DEN induced tumors (Figure 5A, Supplementary Table 9), we decided to further examine this pathway. Of the three serine biosynthesis enzymes, PHGDH is one of the most up-regulated in the DEN tumors (Figure 5A). The enzyme PHGDH shuttles 3-phosphoglycerate from glycolysis and produces serine in collaboration with subsequent reactions catalyzed by PSAT and PSPH (Figure 5B). We verified that this upregulation of Phgdh transcript resulted in higher abundance of PHGDH protein with histological staining for PHGDH and immunoblotting (Figure 5C-D). The vast majority of DEN tumors stained positively for PHGDH.

**Figure 5.**
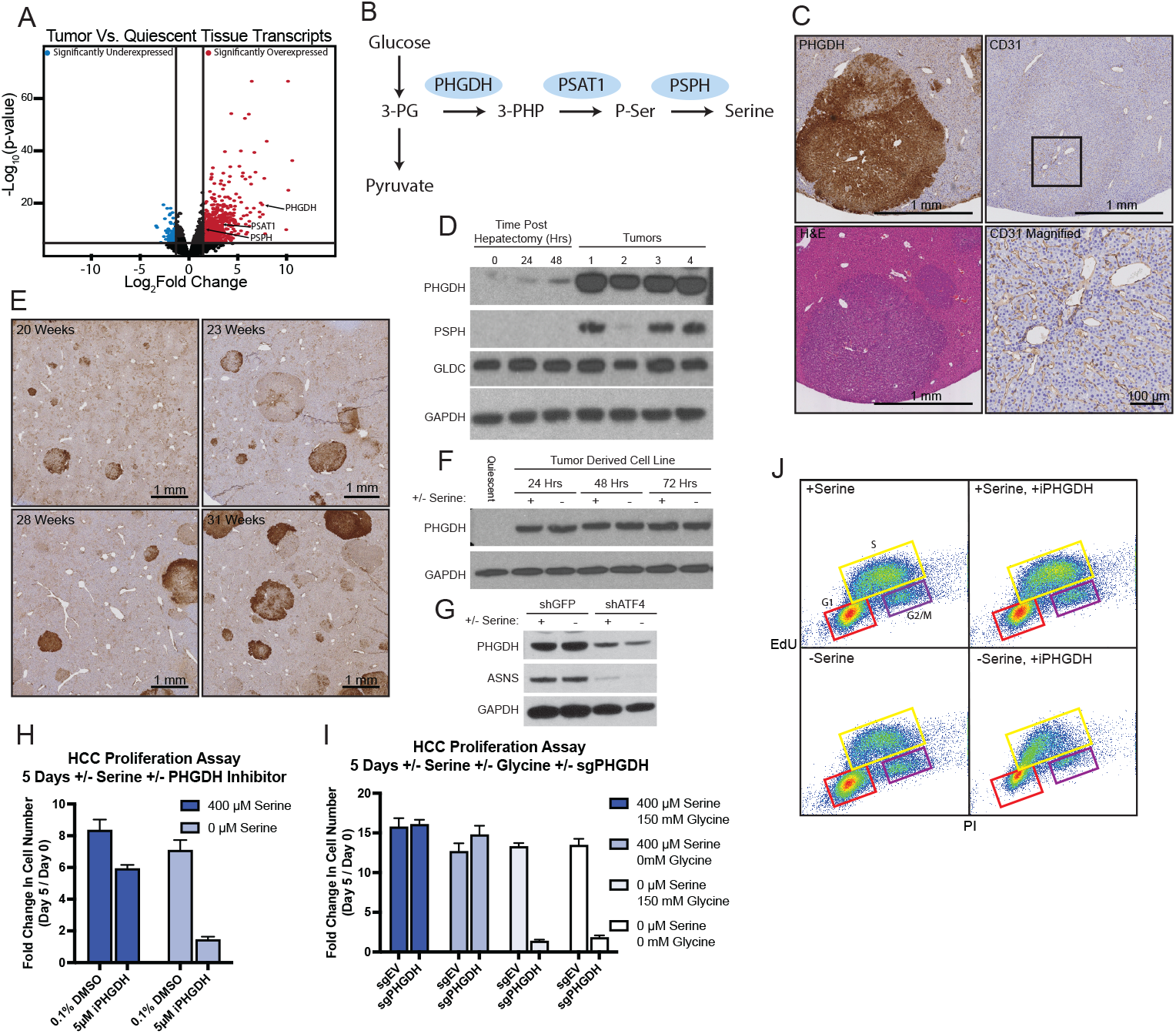
Serine synthesis genes are selectively upregulated in liver transformation. A, Volcano plot of transcripts altered in liver tumors compared to quiescent liver tissue, Log_2_ fold-change >2 and -Log_10_(p) >2 cutoffs indicated. Serine synthesis enzymes indicated. B, Schematic of the serine synthesis pathway. C, Representative immunohistochemical stain (PHGDH, CD31, or hematoxylin and eosin, as indicated) of formalin fixed tissue sections collected from the livers of DEN treated mice 30 weeks after treatment. D, Immunoblot of indicated proteins in lysates from representative tumors versus regenerating tissue. PhX= partial hepatectomy. E, Representative immunohistochemical stain (PHGDH) of formalin fixed tissue sections collected from the livers of DEN treated mice at indicated time point after treatment. F, Immunoblot of indicated proteins from lysates of mouse liver tumor cell line grown in media with or without serine for the indicated lengths of time. G, Immunoblot of indicated proteins from lysates of mouse liver tumor cells infected with a control shRNA (shGFP) or shRNA targeting ATF4 (shATF4). H-I, Fold change in cell number of mouse liver tumor cell lines grown in complete media or media lacking serine and/or glycine as indicated, and upon treatment with vehicle or a PHGDH inhibitor (PH-755, 5 μM). sgEV = empty vector. sgPHGDH = small guide targeting PHGDH. J, EdU incorporation and PI staining of cells treated as in H after 48 hours.

We then determined at what point during tumor development PHGDH protein abundance elevates above the levels in the quiescent tissue. We performed immunohistochemical staining for PHGDH in a set of tissue samples collected at several time points post DEN treatment (Figure 5E). We found strong PHGDH staining by 20 weeks, the earliest time in which we were able to visualize tumors. PHGDH protein abundance was elevated in lesions smaller than one-quarter millimeter in diameter. From this information, we deduce that the acquisition of PHGDH expression is a characteristic obtained at or near the time of transformation.

To assess serine biosynthesis, we generated a cell line from one of these tumors. PHGDH expression was insensitive to exogenous serine treatment (Figure 5F), but dependent upon expression of the integrated stress response transcription factor ATF4 (Figure 5G), as previously described in other systems.^24^ Interestingly, other known ATF4 targets were not upregulated in DEN-induced tumors, indicating that another factor exists which specifically upregulates serine synthesis in DEN-induced transformation. Inhibition of PHGDH with a small molecule inhibitor (PH-755) had minimal effect on cells grown in medium containing serine, while growth in media depleted of serine was impaired, consistent with functional serine biosynthesis (Figure 5H). We further demonstrated pathway functionality by generating a PHGDH knockout cell line using CRISPR/Cas9-mediated gene disruption. PHGDH null cells grown in serine depleted medium exhibited the same sensitivity as the wild type cells treated with the inhibitor, an effect which was not rescued by glycine supplementation (Figure 5I), consistent with prior observations in other systems.^25,26^ To ascertain whether these cells stall during the cell cycle, we performed a dual propidium iodide (PI) / EdU incorporation flow cytometry assay. Tumor cells grown in media lacking serine and treated with the PHGDH inhibitor exhibit cell cycle arrest at early S/G1 phase, consistent with the known requirement for serine-derived carbon in the synthesis of nucleotides (Figure 5J).^26^ These data demonstrate that DEN-induced tumor cells functionally activate serine synthesis, which can support cell cycle progression in the absence of serine.

## Discussion

Many cancer therapies broadly affect both transformed and healthy tissues, highlighting the importance in identifying drug targets specific to tumors. Most studies aimed at solving this problem are either carried out in vitro and lacking strong comparisons to the full scope of healthy cell cycle progression. These details necessitate a model where healthy and dividing tissue can be studied over time and compared to tumors of the same tissue. Previous studies comparing the regenerating liver and transformed liver tissue using partial hepatectomy include regenerative tissue collected at a single time point 24 hours post hepatectomy which, while valuable, does not capture the full spectrum of changes afforded by assaying several time points throughout the regenerative process.^27^

Here we describe our approach to identify metabolic pathways preferentially altered during carcinogen induced tumorigenesis. We identified metabolic pathways known to be altered in cancer as specific for the transformed state in our models, including glycolysis, polyamine catabolism, and serine biosynthesis. Alternatively, healthy regenerating tissue shows preferential activation in glutathione synthesis, triglyceride metabolism, and methionine metabolism.

Serine metabolism has recently been gained focus as an important pathway rewired in cancer.^28,29^ Cancer cells undergoing uncontrolled proliferation require an abundance of serine, either through uptake from the environment or through de novo biosynthesis.^26,30,31^ Cancers that exhibit elevated serine biosynthesis, including some hepatocellular carcinoma, lung adenocarcinoma and breast cancers, through up regulation of the biosynthetic enzymes PHGDH, PSAT1, and PSPH generally show poorer prognosis.^25,28^ Multiple hypotheses have been put forward for why serine synthesis is elevated in the transformed state, including supplying serine in times of high demand, providing TCA cycle intermediates through the action of the transaminase PSAT1, surviving in serine-poor environments, or production of the oncometabolite 2-hydroxyglutarate.^25,32-34^ Therapies to target altered serine biosynthesis in cancer include inhibitors of PHGDH and dietary serine restriction, which can limit tumor growth in a variety of models,^35-39^ while others exhibit resistance despite elevated PHGDH or serine synthesis.^35,40^ The data presented here indicate that PHGDH activation is a common and very early event in RAS/RAF-driven tumorigenesis, and that its activation is not simply associated with cell cycle progression or proliferation.

A subset of the changes we observed in the regenerating liver that are not observed in liver tumors, including altered triglyceride metabolism, may be specifically required for maintaining liver function during regeneration rather than general cell cycle progression. The transient steatosis present during regeneration is consistent with a metabolic program that acquires triglycerides from the circulation and/or retains them from the diet via the portal circulation, to support the regenerative process. This accumulation of lipids in the liver is likely distinct from that present in non-alcoholic fatty liver disease (driven largely by chronic changes in insulin and glucose levels^41^), alcoholic fatty liver disease (driven by inhibition of fatty acid oxidation and lipid handling by ethanol^42^) or fasting-induced steatosis (driven by lipolysis in adipose tissue and import into the liver for oxidation^43^).

Induction of carcinogenesis through DEN administration leads to tumors with mutations in Braf, Hras, and Egfr. These mutations are less common in human hepatocellular carcinoma, which often present activating mutations in beta-catenin. However, the small subset of human HCC containing activating mutations in HRAS and BRAF often lead to more aggressive tumors with poorer prognosis.^44^ Therefore, the model presented here may be representative of an aggressive subset of human HCC.

## Acknowledgements

We thank Possemato lab members for helpful comments. Research is supported by the NIH (T32CA009161, CA168940, GM132491, P30CA016087). R.P. is a scholar of the Pew Charitable Trusts and the Alexander and Margaret Stewart Trust, and an American Cancer Society Research Scholar. plentiCRISPR v2 was a gift from Feng Zhang. PH-755 provided by Michael Pacold.

## Author Contributions

Conceptualization, R.P., D.J.M..; Methodology, R.P., D.J.M.; Investigation, D.J.M, M.N., H.M.; Writing – Original Draft, D.J.M. and R.P.; Writing – Review & Editing, Funding Acquisition, Resources, and Supervision, R.P.

## SUPPLEMENTARY FIGURES

**Supplementary Figure 1.**
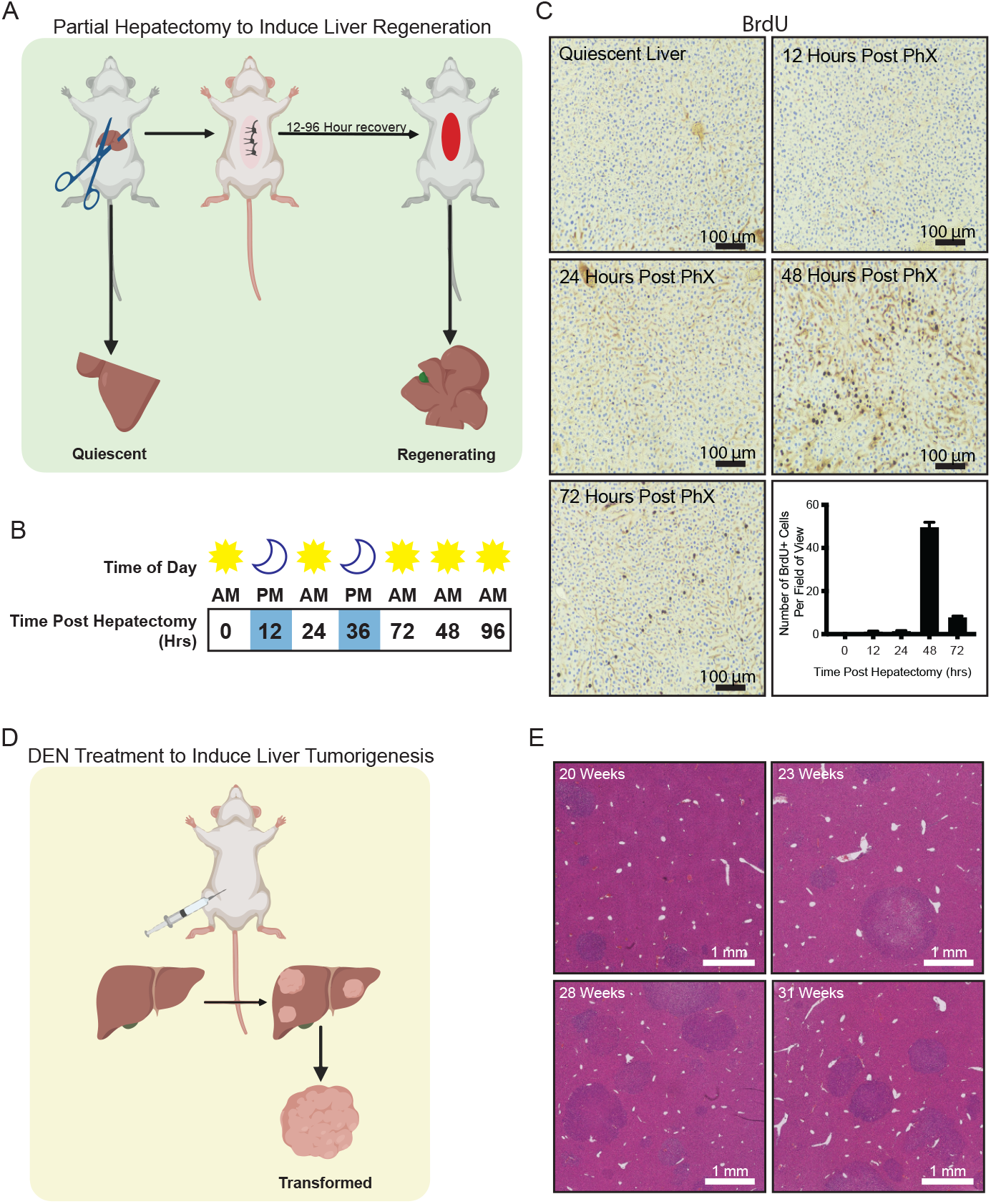
Experimental Set-up for Liver Regeneration and Tumor Initiation Models. A, Experimental design for partial hepatectomy induced liver regeneration. Tissue collected at indicated time points for RNA and metabolite isolation. B, Schematic showing time of tissue collection. Tissue was collected at 10-11am or 10-11pm. C, Immunohistochemical staining and quantification of BrdU incorporation, representative images shown. Bar graph reports the average number of positive cells at each time point (n=5, error bars are s.e.m.). D, Experimental design for tumor initiation. Tissue collected at 30 weeks for RNA and metabolite isolation. E, Hematoxylin and eosin stain of microscopic liver tumors at indicated time after DEN treatment.

**Supplementary Figure 2.**
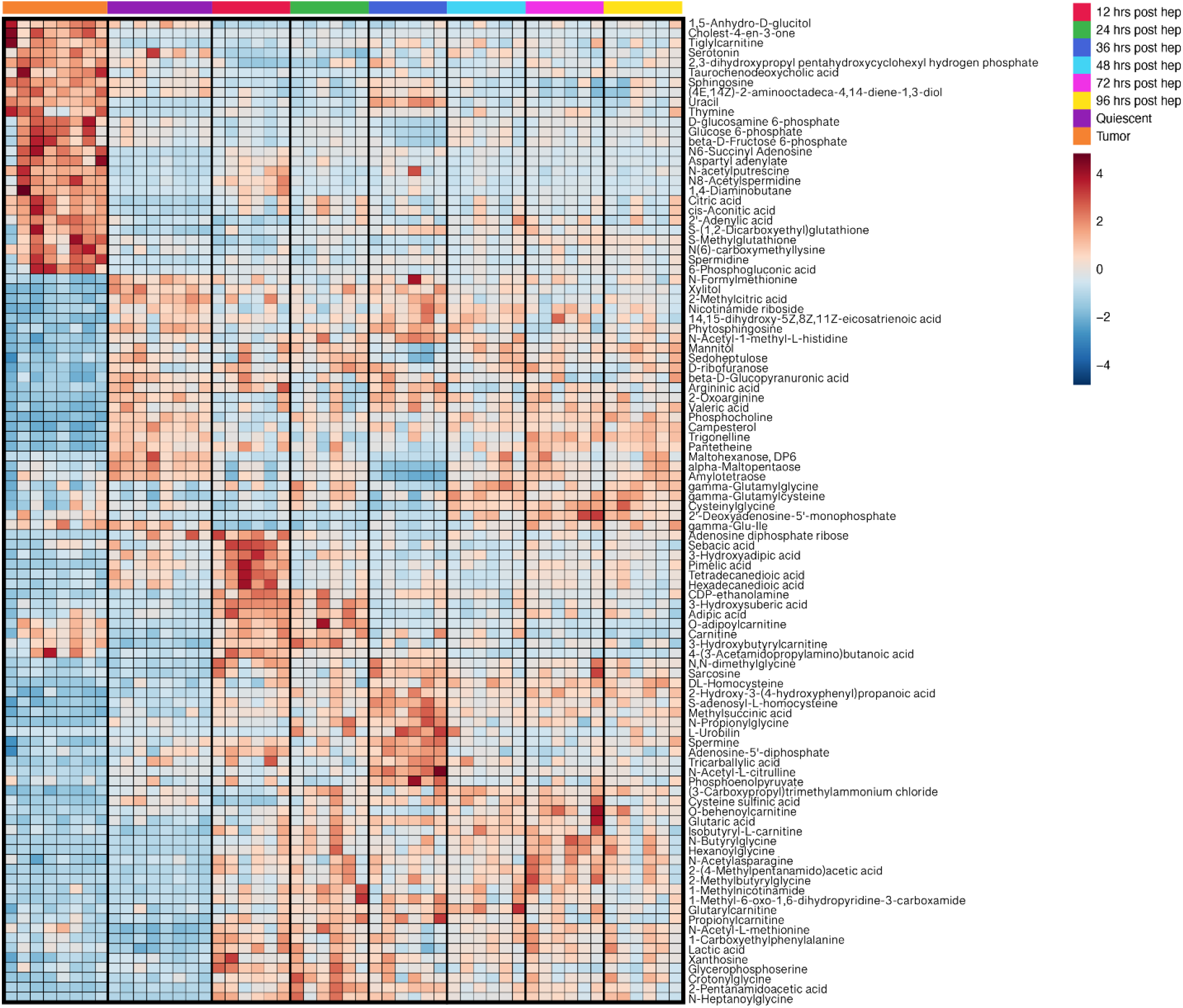
Significantly altered metabolites during regeneration or transformation. Hierarchically clustered heatmap of significantly altered metabolites in both DEN-induced liver tumors, regenerating liver tissue, and quiescent liver samples, p < 0.05 (ANOVA) for any group (regeneration or tumor) compared to quiescent tissue. Relative Log_2_ fold change is reported in the heatmap.

**Supplementary Figure 3.**
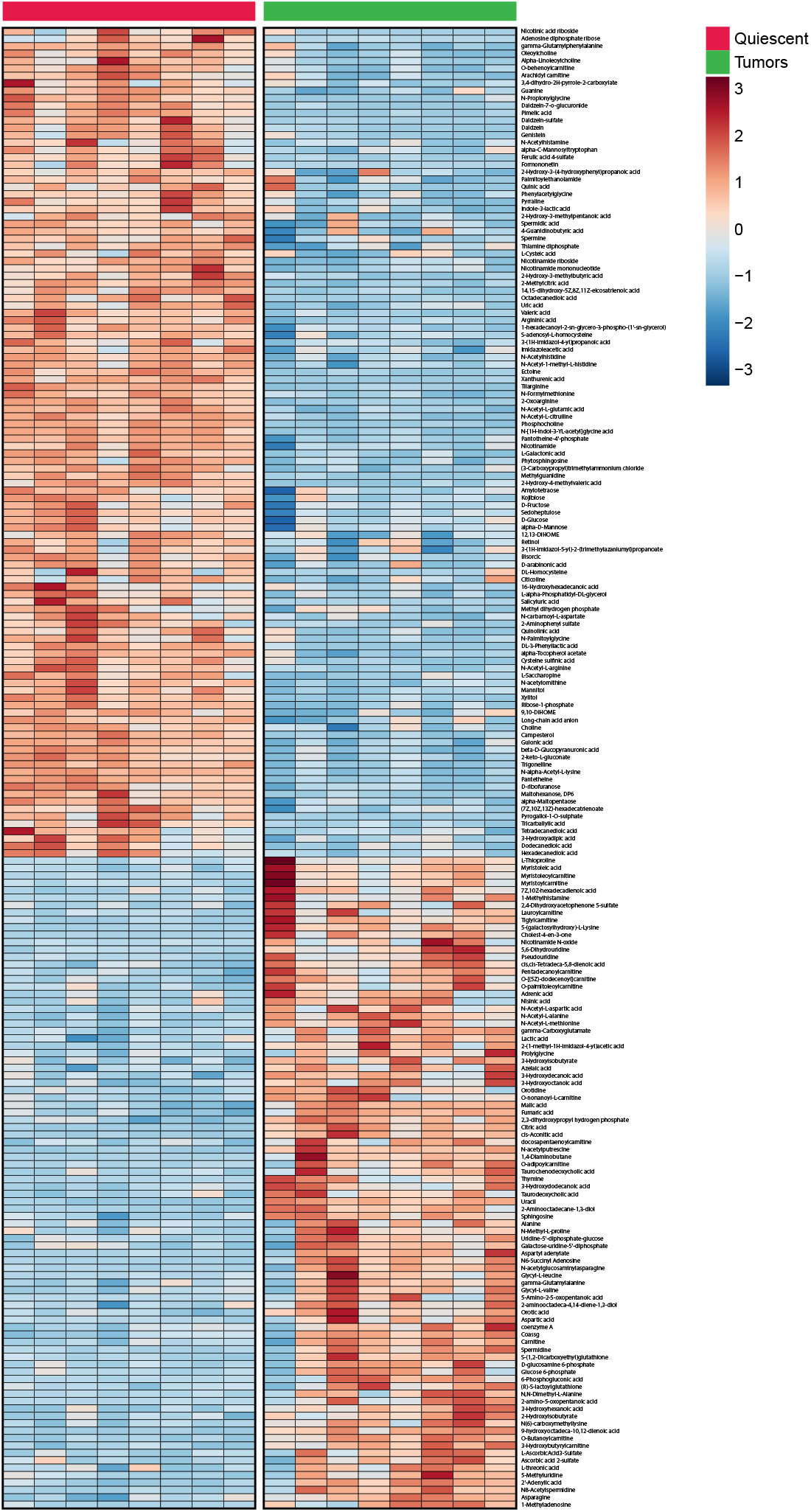
Significantly altered metabolites upon transformation. Hierarchically clustered heatmap of significantly altered metabolites in DEN-induced liver tumors versus quiescent liver samples, p <0.05. Relative Log_2_ fold change is reported in the heatmap.

**Supplementary Figure 4.**
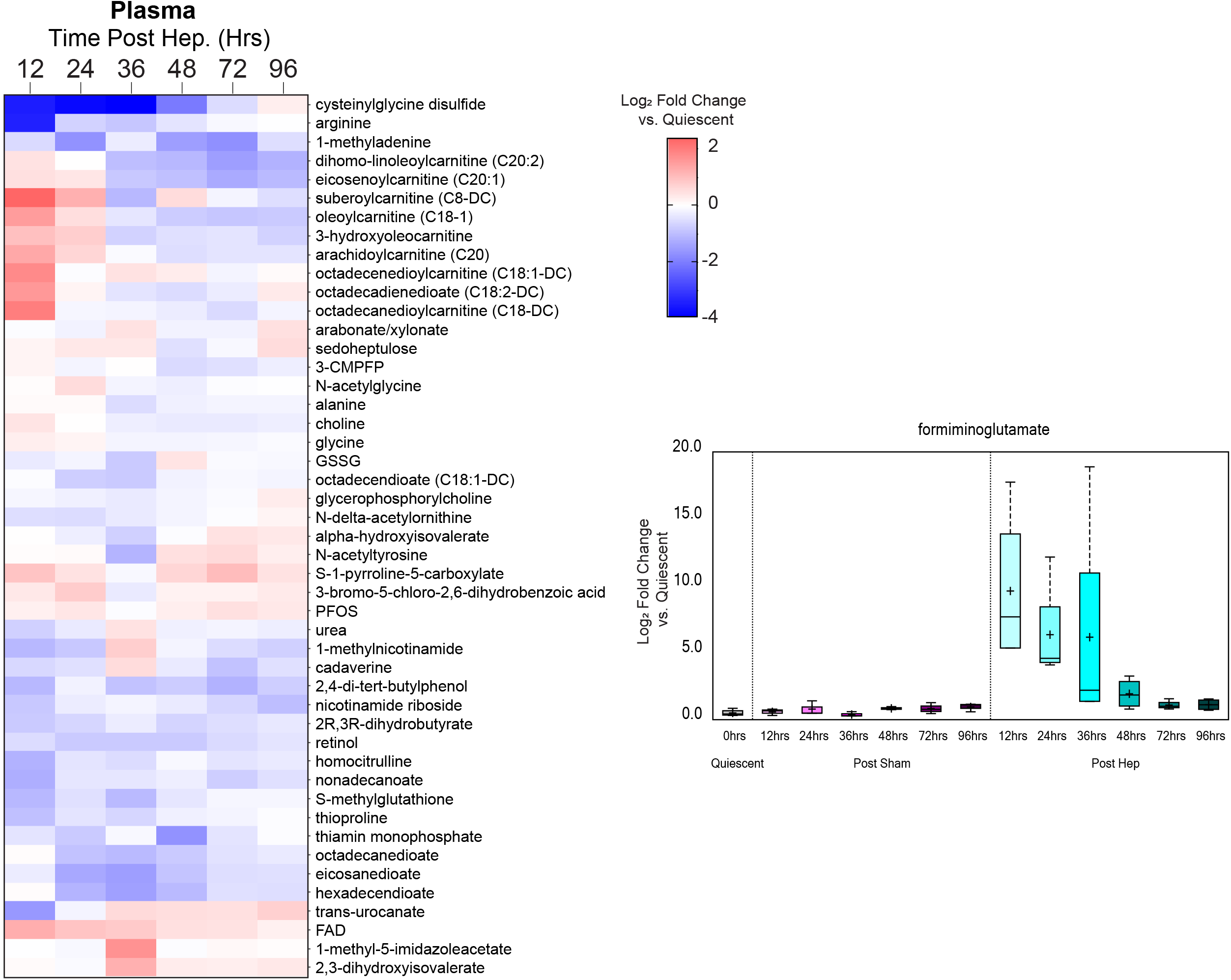
Significantly altered metabolites in plasma during regeneration. Hierarchically clustered heatmap of significantly altered metabolites from plasma at the indicated time point after partial hepatectomy (p < 0.01, quiescent, n=8; 12-96 hr. regeneration time points, n=6). Log_2_ fold change relative to quiescent reported. Boxplot on right shows data for plasma formiminoglutamate and the indicated time points after sham or partial hepatectomy. Boxplots indicate maximum, upper quartile, median, lower quartile, and minimum. “+” indicates mean.

## SUPPLEMENTARY TABLES

**Supplementary Table 1.RNAseq data for quiescent, regeneration, tumor samples.**Counts reported for each gene and sample.

**Supplementary Table 2.RNAseq data for quiescent and sham samples.**Counts reported for each gene and sample.

**Supplementary Table 3.Relative abundance of metabolites from quiescent, regeneration, tumor tissue samples.**

**Supplementary Table 4.Relative abundance of metabolites from quiescent and regeneration plasma samples.**

**Supplementary Table 5.Concentration of lipid metabolites from quiescent and regeneration tissue samples.**

**Supplementary Table 6.Concentration of lipid metabolites from quiescent and regeneration plasma samples.**

**Supplementary Table 7.Order of metabolites and genes in heat maps from main figures. Supplementary Table 8. Genotype of tumors analyzed for common oncogenes.**

**Supplementary Table 9.Data for volcano plat in Figure 6A.**Liver tumor and quiescent tissue transcript abundance compared.

## METHODS

### Animals

C3HeJ wild type male mice were used for all mouse related experiments. All experiments involving mice were carried out with approval from the Committee for Animal Care and under supervision of the Department of Comparative Medicine at MIT and NYU Langone Medical Center.

### Partial Hepatectomy

Adult mice aged 25-30 weeks old were anesthetized using a cocktail of ketamine / xylazine. For sham surgeries, an incision was made in the peritoneal cavity and the liver was gently manipulated with a sterile cotton swab. The peritoneal cavity was then sutured closed, and the skin of the mouse was sealed using wound clips. For hepatectomy surgeries, after anesthesia, the peritoneal cavity was opened, and a knot of sterile nylon filament was tied around the left lobe of the liver. The left lobe was removed below the nylon knot to prevent bleeding. The resected quiescent tissue was quickly rinsed in sterile saline and either preserved in formalin or snap frozen in liquid nitrogen. The peritoneal cavity was sutured shut and the skin was sealed using skin staples. After 12, 24, 36, 48, 72, or 96 hours the mouse was euthanized, and the remnant liver was quickly removed and preserved in the same manner as before.

### DEN Tumor formation

14 day old male mice were treated via intraperitoneal cavity injection with 25mg/kg DEN dissolved in sterile saline. starting at 20 weeks post treatment, groups of mice were euthanized, and their livers were collected and preserved in formalin for subsequent histological staining. At 30 weeks post treatment, the majority of mice were euthanized, and their livers were removed. A portion of each liver was preserved in formalin, while Individual tumors were resected from the remaining liver portions. These tumors were rinsed in sterile saline and snap frozen in liquid nitrogen.

### Transcriptomics

30 mg of tissue from quiescent liver, regenerating liver, or tumors were rinsed in sterile saline and homogenized in Triazole according to Qiagen specifications for the RNeasy Plus Universal Mini Kit with the additional step of gDNA elimination. The resulting RNA was submitted to the NYU Genome Technology Center for paired end run RNASeq. Samples were demultiplexed and normalized counts were provided by the NYU Applied Bioinformatics Laboratory. Hierarchical clustering and principle component analysis were performed using the edgeR package in R. Pathway and biological process analyses were conducted using DAVID Bioinformatics Resources online software. Pathways with an FDR < 0.5 were selected for subsequent analysis.

### Metabolite Profiling

Whole tumors were collected, rinsed in saline, and sliced over dry ice into 30 mg to 50 mg pieces. Sample extraction and metabolite profiling using liquid chromatography-mass spectrometry (LC-MS) was provided by Metabolon. Samples were prepared using MicroLab STAR system (Hamilton Company). Recovery standards were added to each sample prior to extraction for quality control. Samples were extracted using methanol and centrifugation. The extract was separated into 4 fractions: 1) and 2) Used for reverse phase Ultra High Performance LC-MS/MS (UPLC-MS/MS) with positive ion mode electrospray ionization. 3) Used for negative ion mode ESI. 4) used for negative ion ESI HILIC/UPLC-MS/MS. TurboVap (Zymark) was used to remove organic solvent from the samples. The following controls were used in the preparation of metabolite samples: Pooled matrix of all samples, an aliquot of ultra-pure water as a process blank, an aliquot of solvent used for extraction as a solvent blank, and a pool of previously characterized human plasma as a method to determine protocol viability. Recovery standards were used to assess variability of the extraction process. Internal standards were used to assess variability in instrument performance. Metabolon’s proprietary hardware and software were used to extract raw data, identify peaks, and assess quality control. Metabolite identification was based on retention tume index, mass to charge ratio, and MS/MS forward and reverse scores between experimental data and controls. Area-under-the-curve was used to quantify and normalize metabolite peaks. Pathway analysis, statistical calculations, and hierarchical clustering were performed using the MetaboAnalyst package of software.

### Immunohistochemistry

Immunohistochemistry was conducted by the NYU Experimental Pathology Core. For H&E, PHGDH, and CD31 staining, liver tissue samples were preserved in 10% formalin for 48 hours and subsequently stored in 70% ethanol until processed by the core. For Oil Red O staining, samples were preserved in Richard-Allan Scientific “NEG-50” gel and slowly frozen in plastic cassettes over a dry ice ethanol bath. These samples were sliced and stained by the NYU Experimental Pathology Core.

### Cell Culture

Cells were tested to be mycoplasma free by PCR based methods. Cells were cultured in DMEM/F12 media supplemented with 10% IFS (Sigma) and penicillin/streptomycin unless otherwise noted.

### DEN Tumor Cell Line Isolation

30 weeks after DEN administration, mice were euthanized. Tumors were removed, rinsed in cold PBS, and transferred to a sterile petri dish in a tissue culture hood. The tumors were minced using sterile iris scissors and incubated in serum free DMEM/F12 with 1mg/mL collagenase for 2 hours rocking at 37 degrees Celsius. The mixture was centrifuged at 200xg for 3 minutes and washed 3 times in PBS. The pellet was resuspended in 2ml Trypsin + EDTA and incubated for 5-15 minutes at 37 degrees Celsius rocking. The mix was centrifuged for 3 minutes at 200xg, washed 3 times in PBS and resuspended in DMEM. The media was changed every day for 3 days.

### Generation of Knockout Cell Lines

The following primers were selected to make small guides for PHGDH knockout experiments: SgPHGDH-1

5’caccgAGGTGCTCCCTACCAAGCCG

5’aaacCGGCTTGGTAGGGAGCACCTc SgPHGDH-2 5’caccgTGAGCCCGGAATACGAGCAG

5’aaacCTGCTCGTATTCCGGGCTCAc

SgRNAs were cloned into pLentiCRISPR-v2 linearized with BsmBI with Quick ligase (New England Biolabs; M2200). These vectors along with lentiviral packaging vectors Delta-VPR and CMV VSV-G were transfected into HEK293FT cells by polyelthylenimine (Polysciences, Inc; 60 μg/mL) mediated transfection. Media was changed 24 hr after transfection and the virus-containing supernatant was collected 48 hr after transfection. Virus was passed through a 0.45 μm filter and stored at -80 °C or used immediately. Target cells in 6-well tissue culture plates were infected in media containing 2 μg/ml polybrene by spin infection at 2,250 rpm for 30 min. 24 hours post-infection, virus was removed and cells were selected with puromycin. After selection single cells were plated into the wells of a 96-well plate. Cells were grown for 2-3 weeks, and the resultant colonies were expanded and screened for loss of the relevant protein by immunoblotting. SgPHGDH-1 was chosen for subsequent assays.

### Immunoblotting

Tumors were rinsed in ice-cold saline and sliced over dry ice into fragments ranging in size from 15 to 30 mg. each fragment was homogenized using a Precelyss 24 homogenizer at 6000 rpm for 20 seconds or until the tissue was dissolved. The tumors were homogenized in lysis buffer containing 50 mM Hepes, pH 7.4, 40 mM NaCl, 2 mM EDTA, 50 mM NaF, 10 mM pyrophosphate, 10 mM glycerophosphate, protease inhibitors (Roche) and 1% Triton-X-100. After homogenization, the samples were centrifuged at 4 degrees Celsius for 10 minutes at maximum speed. Supernatant was transferred to new tubes and Protein concentrations were measured by the Bradford method and 15 μg of total lysates were resolved by SDS-PAGE (4-12% gel) and analyzed by immunoblotting. Cells were rinsed once in ice-cold PBS and harvested in a lysis buffer containing 50 mM Hepes, pH 7.4, 40 mM NaCl, 2 mM EDTA, 50 mM NaF, 10 mM pyrophosphate, 10 mM glycerophosphate, protease inhibitors (Roche) and 1% Triton-X-100. Protein concentrations were measured by the Bradford method and 15 μg of total lysates were resolved by SDS-PAGE (4-12% gel) and analyzed by immunoblotting. Sample gels were transferred at 0.3 constant amps for 90 minutes on PVDF membranes. Membranes were blocked using 5% nonfat milk for 30 minutes and incubated overnight in 5% milk with the following antibodies: PHGDH (1:1000, Sigma HPA021241), GAPDH (1:5000, CST 5174), PSPH (1:1000, Sigma HPA020376), and GLDC (1:1000, CST 12794S. The membranes were then washed in 5% TBS-T for 5 minutes 3 times and incubated with the following secondary antibody for 1 hour at room temperature in 5% milk: Goat anti-Rabbit IgG (H+L) HRP (1:5000, CST 31460). Membranes were imaged using Pierce ECL western blotting substrate. Optimal exposure was determined to be between 5 seconds and 5 minutes for each target.

### Proliferation Assays

Direct cell counts were carried out by plating cells in triplicate in 12-well plates at 10,000 cells per well in 2 mL of DMEM F/12 containing 10% dialyzed FBS and any combination of 0 µM serine, 400 µM serine, 0 µM glycine, or 150 mM glycine. After 5 days, cells were counted using a Beckman Z2 Coulter Counter with a size selection setting of 8 to 30 μm.

### Propidium Iodide and EdU Flow Analysis

Assay was performed using EdU Cytometry Alexa Fluor 647 kit from Invitrogen (C10419, C10420). 10 µM EdU was added to the culture medium and incubated for 2 hours. Cells were washed in PBS and harvested using trypsin. Cell suspension was washed in PBS containing 1% BSA and centrifuged at 500xg for 5 minutes. The cell pellet was resuspended in 100 µL Click-iT fixative and mixed well. The mixture was then incubated for 15 minutes in a dark environment. The cells were then washed with 3 mL of PBS with 1% BSA and centrifuged for 5 minutes at 400xg. The pellet was resuspended in 100 µL saponin based permeabilization and wash reagent and incubated for 15 minutes. Click-iT reaction mix was prepared according to kit protocols. 500 µL of reaction mix were added to each reaction tube and incubated for 30 minutes in a dark environment. Samples were washed in 3 mL permeabilization buffer, centrifuged at 500xg for 5 minutes, and resuspended in permeabilization buffer with Ribonuclease A added. 10 µL of 0.5 mg/mL Propidium Iodide were added to each tube. Cell samples were analyzed in an Attune NxT flow cytometer. EdU and Propidium Iodide were analyzed using the red (638nm) and blue (488 nm) lasers respectively. Results were interpreted using FlowJo software.

### Figures

Graphical abstracts and summary figures were created using Biorender online software. All other figures created using Adobe Illustrator.

### Data availability

The data generated in this study are available within the article and its Supplementary data files.

### Quantification and Statistical Analysis

No samples or animals were excluded from analysis and sample size estimates were not used. The number of independent biological replicates (n) are indicated in the figure legend and represent replicate measurements from distinct samples. For immunoblots, the reported images are representative of at least three independent experiments. Studies were not conducted blind. DAVID software used for analysis of significantly enriched genes sets is at https://david.ncifcrf.gov/

